# Multicenter validation of a sepsis prediction algorithm using only vital sign data in the emergency department, general ward and ICU

**DOI:** 10.1101/243964

**Authors:** Qingqing Mao, Melissa Jay, Jana L. Hoffman, Jacob Calvert, Christopher Barton, David Shimabukuro, Lisa Shieh, Uli Chettipally, Grant Fletcher, Yaniv Kerem, Yifan Zhou, Ritankar Das

**Author notes:** Corresponding author: 22710 Foothill Blvd., Suite #2 Hayward, CA 94541.

## Abstract

**Objectives:** We validate a machine learning-based sepsis prediction algorithm (*InSight*) for detection and prediction of three sepsis-related gold standards, using only six vital signs. We evaluate robustness to missing data, customization to site-specific data using transfer learning, and generalizability to new settings.

**Design:** A machine learning algorithm with gradient tree boosting. Features for prediction were created from combinations of only six vital sign measurements and their changes over time.

**Setting:** A mixed-ward retrospective data set from the University of California, San Francisco (UCSF) Medical Center (San Francisco, CA) as the primary source, an intensive care unit data set from the Beth Israel Deaconess Medical Center (Boston, MA) as a transfer learning source, and four additional institutions’ datasets to evaluate generalizability.

**Participants:** 684,443 total encounters, with 90,353 encounters from June 2011 to March 2016 at UCSF.

**Interventions:** none

**Primary and secondary outcome measures:** Area under the receiver operating characteristic curve (AUROC) for detection and prediction of sepsis, severe sepsis, and septic shock.

**Results:** For detection of sepsis and severe sepsis, *InSight* achieves an area under the receiver operating characteristic (AUROC) curve of 0.92 (95% CI 0.90 - 0.93) and 0.87 (95% CI 0.86 - 0.88), respectively. Four hours before onset, *InSight* predicts septic shock with an AUROC of 0.96 (95% CI 0.94 -0.98), and severe sepsis with an AUROC of 0.85 (95% CI 0.79 - 0.91).

**Conclusions:** *InSight* outperforms existing sepsis scoring systems in identifying and predicting sepsis, severe sepsis, and septic shock. This is the first sepsis screening system to exceed an AUROC of 0.90 using only vital sign inputs. *InSight* is robust to missing data, can be customized to novel hospital data using a small fraction of site data, and retained strong discrimination across all institutions.

**Strengths and limitations of this study:**

- Machine learning is applied to the detection and prediction of three separate sepsis standards in the emergency department, general ward and intensive care settings.
- Only six commonly measured vital signs are used as input for the algorithm.
- The algorithm is robust to randomly missing data.
- Transfer learning successfully leverages large dataset information to a target dataset.
- Retrospective nature of the study does not predict clinician reaction to information.

## Introduction

Sepsis is a major health crisis and one of the leading causes of death in the United States [1]. Approximately 750,000 hospitalized patients are diagnosed with severe sepsis in the United States annually, with an estimated mortality rate of up to one-third [2,3].The cost burden of sepsis is disproportionately high, with estimated costs of $20.3 billion dollars annually, or $55.6 million per day in US hospitals [4]. Additionally, the average hospital stay for sepsis is twice as expensive as other conditions [5], and the average incidence of severe sepsis is increasing by approximately 13% per year [6]. Early diagnosis and treatment have been shown to reduce mortality and associated costs [7–9]. Despite clear benefits, early and accurate sepsis detection remains a difficult clinical problem.

Sepsis has been defined as a dysregulated host response to infection. In practice, sepsis can be challenging to recognize because of the heterogeneity of the host response to infection, and the diversity of possible infectious insult. Sepsis has been traditionally recognized as two or more Systemic Inflammatory Response Syndrome (SIRS) [10] criteria together with a known or suspected infection; progressing to severe sepsis, in the event of organ dysfunction; and finally to septic shock, which additionally includes refractory hypotension [10]. However, ongoing debates over sepsis definitions and clinical criteria, as evidenced by the recent proposed redefinitions of sepsis [11], underscore a fundamental difficulty in the identification and accurate diagnosis of sepsis.

Various rule-based disease severity scoring systems are widely used in hospitals in an attempt to identify septic patients. These scores, such as the Modified Early Warning Score (MEWS) [12], the Systemic Inflammatory Response Syndrome (SIRS) criteria [13], and the Sequential Organ Failure Assessment (SOFA) [14], are manually tabulated at the bedside and lack accuracy in sepsis diagnosis. However, the increasing prevalence of Electronic Health Records (EHR) in clinical settings provides an opportunity for enhanced patient monitoring and increased early detection of sepsis.

This study validates a machine learning algorithm *InSight*, which uses only six vital signs taken directly from the EHR, in the detection and prediction of sepsis, severe sepsis, and septic shock in a mixed-ward population at the University of California, San Francisco (UCSF). We investigate the effects of induced data sparsity on *InSight* performance, and compare all results with other scores that are commonly used in the clinical setting for the detection and prediction of sepsis. We additionally train and test the algorithm for severe sepsis detection on data from Stanford Medical Center and three community hospitals in order to better estimate its expected clinical performance. Furthermore, we apply a transfer learning scheme to customize a Multiparameter Intelligent Monitoring in Intensive Care (MIMIC)-III-trained algorithm to the UCSF patient population using a minimal amount of UCSF-specific data.

## Methods

### Data sets

We used a data set provided by the UCSF Medical Center representing patient stays from June 2011 to March 2016 in all experiments. The UCSF data set contains 17,467,987 hospital encounters, including inpatient and outpatient visits to all units within the UCSF medical system. The data were de-identified to comply with the Health Insurance Portability and Accountability Act (HIPAA) Privacy Rule. For transfer learning, we used the Multiparameter Intelligent Monitoring in Intensive Care (MIMIC)-III v1.3 data set, compiled from the Beth Israel Deaconess Medical Center (BIDMC) in Boston, MA between 2001 and 2012, composed of 61,532 ICU stays [15]. This database is a publicly available database constructed by researchers at MIT’s Laboratory for Computational Physiology, and the data were also de-identified in compliance with HIPAA. Additionally, we trained and tested the algorithm for severe sepsis detection on data from Stanford Medical Center (Stanford, CA), Oroville Hospital (Oroville, CA), Bakersfield Heart Hospital (BHH; Bakersfield, CA), and Cape Regional Medical Center (CRMC; Cape May Courthouse, NJ). Details on these datasets are included in the Supplementary Materials (Supplementary Tables 1 and 2). Data collection for all datasets did not impact patient safety. Therefore, this study constitutes non-human subjects research, which does not require Institutional Review Board approval.

### Data Extraction and Imputation

The data were provided in the form of comma separated value (CSV) files and stored in a PostgreSQL [16] database. Custom SQL queries were written to extract measurements and patient outcomes of interest. The measurement files were then binned by hour for each patient. To be included, patients were required to have at least one of each type of measurement recorded during the encounter. If a patient did not have a measurement in a given hour, the missing measurement was filled in using carry-forward imputation. This imputation method applied the patient’s last measured value to the following hour (a causal procedure). In the case of multiple measurements within an hour, the mean was calculated and used in place of an individual measurement. After the data were processed and imputed in Python [17], they were used to train the *InSight* classifier and test its predictions at sepsis onset and at fixed time points prior to onset.

### Gold Standards

In this study, we tested *InSight’s* performance according to various gold standards (clinical indications). We investigated *InSight’s* ability to predict and detect sepsis, severe sepsis, and septic shock. Further, we compared *InSight’s* performance to SIRS, MEWS, and SOFA, for each of the following gold standards. For training and testing the algorithm, we conservatively identified each septic condition by requiring that the ICD-9 code corresponding to the diagnosis was coded for each positive case, in addition to meeting the clinical requirements for the definition of each septic standard as defined below.

### Sepsis

The sepsis gold standard was determined using the 2001 consensus sepsis definition [10]: “the presence of two or more SIRS criteria paired with a suspicion of infection.” To identify a case as positive for sepsis, we required ICD-9 code 995.91. The onset time was defined as the first time two or more SIRS criteria were met within the same hour. SIRS criteria are defined as:

- heart rate > 90 beats/ min,
- body temperature > 38 □ or < 36 □,
- respiratory rate >20 breaths/min or PaCO_2_ < 32 mmHg, and
- white blood cell count > 12,000 cells/μL or < 4,000 cells/μL. [10]

### Severe Sepsis

The severe sepsis gold standard used the definition of severe sepsis as “organ dysfunction caused by sepsis” which can be represented by one or more of the criteria below, and identified for patients with the severe sepsis ICD-9 code 995.92. We assigned the severe sepsis onset time to be the first instance during which two SIRS criteria as described above and one of the following organ dysfunction criteria were met within the same hour.

- Lactate > 2 mmol/L
- Systolic blood pressure < 90 mmHg
- Urine output < 0.5 mL/kg, over two hours, prior to organ dysfunction after fluid resuscitation
- Creatinine > 2 mg/dL without renal insufficiency or chronic dialysis
- Bilirubin > 2 mg/dL without having liver disease or cirrhosis
- Platelet count < 100,000 μL
- International normalized ratio > 1.5
- PaO2/FiO2 < 200 in addition to pneumonia < 250 with acute kidney injury but without pneumonia

### Septic Shock

We identified as positive cases for septic shock those patients who received the septic shock ICD-9 code 785.52 and additionally demonstrated the following conditions:

- systolic blood pressure of < 90 mmHg, defined as hypotension, for at least 30 minutes, and
- who were resuscitated with ≥ 20 ml/kg over a 24 hour period, or
- who received ≥ 1200 ml in total fluids. [18]

The onset time was defined as the first hour when either the hypotension or fluid resuscitation criterion was met.

## Calculating Comparators

We compared *InSight* predictions for each gold standard to three common patient deterioration scoring systems: SIRS, SOFA, and MEWS. Area under the receiver operating characteristic (AUROC) curve, sensitivity, and specificity were compared across all prediction models. The SIRS criteria, as explained in the sepsis definition, were evaluated independently of the suspicion of infection. To calculate the SOFA score, we collected each patient’s PaO_2_/FiO_2_, Glasgow Coma Score, mean arterial blood pressure or administration of vasopressors, bilirubin level, platelet counts, and creatinine level. Each of the listed measurements is associated with a SOFA score of 1-4, based on severity level, as described by Vincent et al. [14]. After receiving a score for each of the six organ dysfunction categories, the overall SOFA score was computed as the sum of the category scores and used as a comparator to *InSight*. Finally, the MEWS score, which ranges from 0 (normal) to 14 (high risk of deterioration), was determined by tabulating subscores for heart rate, systolic blood pressure, respiratory rate, temperature, and Glasgow Coma Score. We used the subscoring system presented in Fullerton et al. [19] to compute each patient’s MEWS score.

## Measurements and Patient Inclusion

In order to generate *InSight* scores, patient data were analyzed from each of the following six clinical vital sign measurements: systolic blood pressure, diastolic blood pressure, heart rate, respiratory rate, peripheral capillary oxygen saturation (SpO_2_), and temperature. We used only vital signs, which are frequently available and routinely taken in the ICU, ED, and floor units. Patient data were used from the course of a patient’s hospital encounter, regardless of the unit the patient was in when the data were collected.

All patients over the age of 18 were considered for this study. For a given encounter, if the patient was admitted to the hospital from the ED, the start of the ED visit is where the analysis began. Patients in our final data sets were required to have at least one measurement for each of the six vital signs. In order to ensure enough data to accurately characterize sepsis predictions at four hours pre-onset, we further limited the study group to exclude patients whose septic condition onset time was within seven hours after the start of their record, which was either the time of admission to the hospital or the start of their ED visit; the latter was applicable only if the patient was admitted through the ED. A smaller window to sepsis onset time would have resulted in insufficient testing data to make 4-hour prediction possible in some cases, which would inappropriately affect performance metrics such as sensitivity and specificity. Patients with sepsis onset after 2,000 hours post-admission were also excluded, to limit the data analysis matrix size. The final UCSF data set included 90,353 patients (Fig. 1) and the MIMIC-III data set contained 21,604 patients, following the same inclusion criteria. Inclusion criteria and final inclusion numbers for the Stanford, Oroville, BHH, and CRMC datasets are included in Supplementary Table 1.

**Figure 1:**
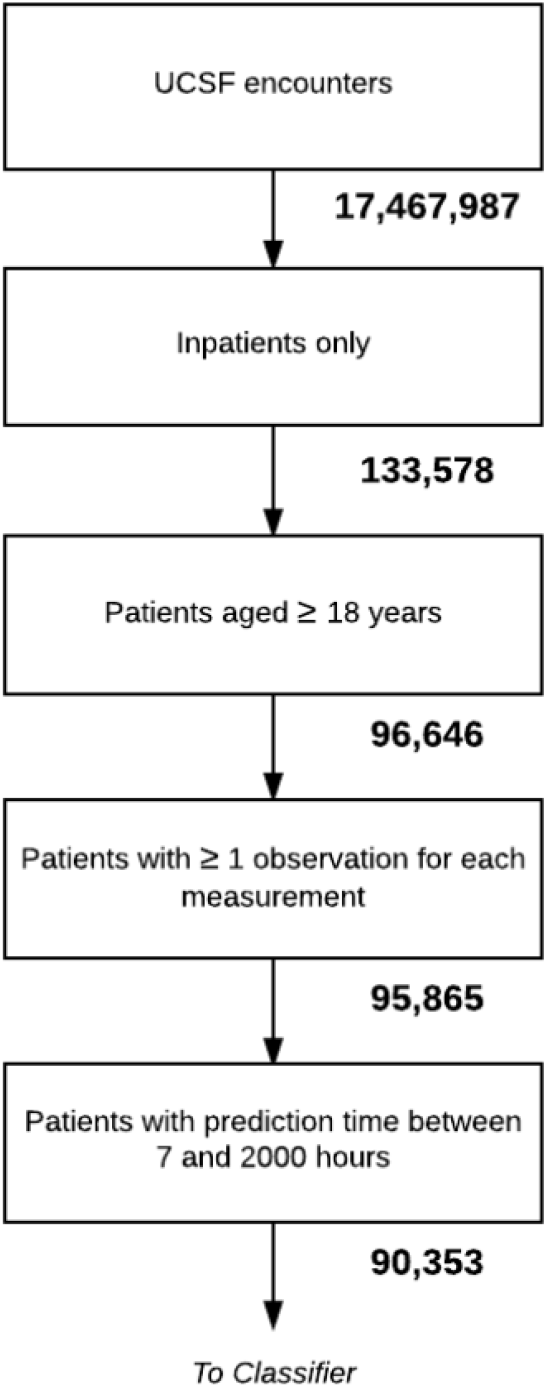
Patient inclusion flow diagram for the UCSF data set.

After patient exclusion, our final group of UCSF patients was composed of 55% women and 45% men with a median age of 55. The median hospital length of stay was 4 days, IQR = (2,6). Of the 90,353 patients, 1,179 were found to have sepsis (1.30%), 349 were identified as having severe sepsis without shock (0.39%), and 614 were determined to have septic shock (0.68%). The in-hospital mortality rate was 1.42%. Patient encounters spanned a variety of wards. The most common units represented in our study were perioperative care, the emergency department, the neurosciences department, and cardiovascular and thoracic transitional care. In the MIMIC-III data set, approximately 44% of patients were women and 56% were men. Stays were typically shorter in this data set, since each encounter included only an ICU stay. The median length of stay was 2 days. Furthermore, due to the nature of intensive care, there was a higher prevalence of sepsis (1.91%), severe sepsis (2.82%), and septic shock (4.36%). A full summary of baseline characteristics for both data sets is presented in Table 1. Full demographic information for the Stanford, Oroville, BHH, and CRMC datasets is provided in Supplementary Table 2.

**Table 1:** Demographic and clinical characteristics for UCSF patient population analyzed (N=90,353) and MIMIC-III patient population analyzed (N=21,604).

| Demographic Overview | Characteristic | UCSF |  | MIMIC-III |  |
| --- | --- | --- | --- | --- | --- |
|  |  | Count | Percentage | Count | Percentage |
| Gender | Female | 49,763 | 55.08% | 9,499 | 43.97% |
|  | Male | 40,590 | 44.92% | 12,105 | 56.03% |
| **Age**<br>UCSF: median 55, IQR (38-67)<br>MIMIC-III: median 65, IQR (53-77) | 18-29 | 10,652 | 11.79% | 978 | 4.53% |
|  | 30-39 | 14,202 | 15.72% | 1,114 | 5.16% |
|  | 40-49 | 11,888 | 13.16% | 2,112 | 9.78% |
|  | 50-59 | 16,856 | 18.66% | 3,880 | 17.96% |
|  | 60-69 | 19,056 | 21.09% | 4,906 | 22.71% |
|  | 70+ | 17,699 | 19.59% | 8,614 | 39.87% |
| **Length of Stay (days)**<br>UCSF: median 4, IQR (2-6)<br>MIMIC-III: median 2, IQR (2-4) | 0-2 | 28,258 | 31.26% | 11,054 | 51.17% |
|  | 3-5 | 35,128 | 38.88% | 7,004 | 32.42% |
|  | 6-8 | 12,664 | 14.02% | 1,673 | 7.74% |
|  | 9-11 | 4,934 | 5.46% | 734 | 3.40% |
|  | 12+ | 9,369 | 10.37% | 1,139 | 5.27% |
| Death During Hospital Stay | Yes | 1,279 | 1.42% | 1,328 | 6.15% |
|  | No | 89,074 | 98.58% | 20,276 | 93.85% |
| ICD-9 Code | Sepsis | 1,179 | 1.30% | 413 | 1.91% |
|  | Severe Sepsis | 349 | 0.39% | 609 | 2.82% |
|  | Septic Shock | 614 | 0.68% | 943 | 4.36% |

## Feature Construction

We minimally processed raw vital sign data to generate features. Following EHR data extraction and imputation as described above, we obtained three hourly values for each of the six vital sign measurement channels from that hour, the hour prior, and two hours prior. We also calculated two difference values between the current hour and the prior hour, and between the prior hour and the hour before that. We concatenated these five values from each vital sign into a causal feature vector ***x*** with 30 elements (five values from each of six measurement channels).

## Machine Learning

We used gradient tree boosting to construct our classifier. Gradient tree boosting is an ensemble technique which combines the results from multiple weak decision trees in an iterative fashion. Each decision tree, was built by discretizing features into two categories. For example, one node of the decision tree might have stratified a patient based on whether their respiratory rate was greater than 20 breaths per minute, or not. Depending on the answer for a given patient, a second, third, etc., vital sign may be checked. A risk score was generated for the patient based on their path along the decision tree. We limited each tree to split no more than six times; no more than 1000 trees were aggregated in the iteration through gradient boosting to generate a robust risk score. Training was performed separately for each distinct task and prediction window, and observations were accordingly labeled positive for model fitting for each specific prediction task. Patient measurements were not used after the onset of a positive clinical indication.

We performed ten-fold cross validation to validate *InSight’s* performance and minimize potential model overfit. We randomly split the UCSF data set into a training set, comprised of 80% of UCSF’s encounters, and an independent test set with the remaining 20% of encounters. Of the training set, data were divided into ten groups, nine of which were used to train *InSight*, and one of which was used to test. After cycling through all combinations of train and test set, we then tested each of the ten models on the independent test set. Mean performance metrics were calculated based on these ten models. For severe sepsis detection at time of onset on each of Stanford, Oroville, BHH, and CRMC datasets, we performed four-fold cross validation of the model.

Additionally, we trained and validated *InSight’s* performance in identifying sepsis, severe sepsis, and septic shock after removing all features which were used in our gold standard definitions for each condition. This resulted in the removal of vital sign SIRS criteria measurements for sepsis and severe sepsis predictions, and the removal of systolic and diastolic blood pressure measurements for septic shock. We also trained and validated the algorithm for each of the three gold standards for randomly selected, up- and down-sampled subpopulations with positive class prevalence between zero and one hundred percent.

## Missing Data

After assessing *InSight’s* performance on complete data sets, we used a random deletion process to simulate the algorithm’s robustness to missing measurements. Individual measurements from the test set were deleted according to a probability of deletion, *P*. We set *P* = {0, 0.1, 0.2, 0.4, and 0.6} for each of our missing data experiments and tested the *InSight* algorithm on the sparse data sets.

## Transfer Learning

To evaluate *InSight’s* performance on a minimal amount of UCSF data, we used a transfer learning approach [20]. There are clear dissimilarities in patient demographics, clinical characteristics, and average measurement frequencies between the UCSF and MIMIC-III data sets (see Table 1). Partially this is because the UCSF data involves a variety of hospital wards, whereas the MIMIC-III data set provides only measurements taken in the ICU. We sought to determine improved performance metrics on the UCSF target data set, when the algorithm is primarily trained on MIMIC-III. Using MIMIC-III data as the source, and UCSF as the target, we trained the *InSight* classifier according to the severe sepsis gold standard. Variable amounts of UCSF training data were incrementally added to the MIMIC-III training data set, and the resulting model was then validated on the separate UCSF test data set. Specifically, we left 50% of the UCSF patients as test data, and we randomly selected different fractions of the remaining UCSF data and combined them with the entire MIMIC-III data set as the training data. For each fraction used, we trained 100 models with different random relative weights on the UCSF and MIMIC-III training data. Then, the mean and standard deviation of AUROC values for each of these models were calculated on 20 randomly sampled sets, and the model with highest mean AUROC value among these 100 was used.

## Results

*InSight’s* performance with respect to MEWS, SOFA, and SIRS is summarized in Figures 2A-C. Figures 2A, 2B, and 2C demonstrate *InSight’s* ability to accurately detect the onset of sepsis and severe sepsis, and to accurately predict septic shock four hours prior to onset, compared to the performance of common sepsis scoring systems. Each figure presents *InSight’s* receiver operating characteristic (ROC) curve together with the ROC curves for MEWS, SOFA, and SIRS. *InSight* achieves an area under the receiver operating characteristic (AUROC) curve for sepsis onset of 0.92 (95% confidence interval (CI), 0.90 - 0.93), for severe sepsis onset of 0.87 (95% CI 0.86 - 0.88), and for septic shock of 0.99 (95% CI 0.9991 - 0.9994); compared to SIRS, which demonstrates an AUROC of 0.75, 0.72, and 0.84, respectively. Even when all gold standard involved measurements were removed from model training, *InSight* continued to demonstrate improved accuracy over SIRS, MEWS, and SOFA, with AUROC values of 0.84 (95% CI 0.83-0.85) for sepsis onset, 0.80 (95% CI 0.79-0.81) for severe sepsis onset, and 0.96 (95% CI 0.96-0.97) for septic shock onset.

**Figure 2:**
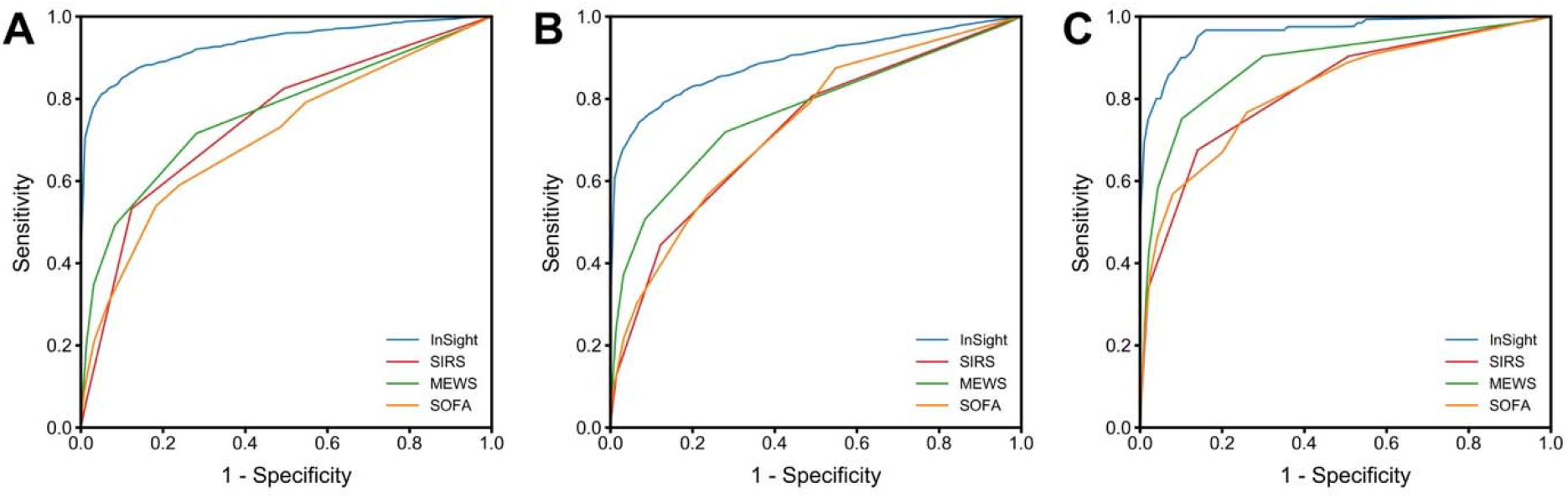
ROC curves for *InSight* and common scoring systems at time of (A) sepsis onset, (B) severe sepsis onset, and (C) four hours before septic shock onset.

Comparing *InSight*’s performance across the three sepsis-related gold standards, it is clear that the septic shock criteria are relatively less challenging to anticipate, as its four hour prediction metrics are stronger than those for the detection of both sepsis and severe sepsis. Accordingly, we display the four hour prior to onset prediction case for septic shock (Fig 2C), where existing tools fail to adequately meet prediction standards relevant for sound clinical use. Four hours in advance of septic shock onset, *InSight* achieved an AUROC of 0.96 (95% CI 0.94 - 0.98). The resulting confusion matrix from the ten-fold cross validation of *InSight* can be found in Supplementary Tables 3 and 4.

Additional comparison metrics at time of detection for each gold standard are available in Table 2. In order to compare the specificities from each gold standard, we fixed sensitivities near 0.80; that is, we fixed a point on the ROC curve (i.e. set a specific threshold) after model development and tested algorithm performance under the chosen conditions in order to present data as consistently as possible. We similarly fixed specificities near 0.80 in order to compare sensitivities. Across all gold standards, a sensitivity of 0.80 results in a high specificity for *InSight*; however, the sensitivities for MEWS, SOFA, and SIRS are significantly lower. Notably, at 0.80 sensitivity, *InSight* achieves a specificity of 0.95 for sepsis, 0.84 for severe sepsis, and 0.99 for septic shock detection.

**Table 2:** Performance metrics for three sepsis gold standards at time of onset (zero hour), with sensitivities fixed at or near 0.80 in the first instance, and specificities fixed at or near 0.80 in the second instance.

|  | Gold Standard | *InSight*<br>(95% CI) | *InSight*,<br>label definitions<br>removed<br>(95% CI) | MEWS | SOFA | SIRS |
| --- | --- | --- | --- | --- | --- | --- |
| AUROC | Sepsis | 0.92<br>(0.90, 0.93) | 0.84<br>(0.83, 0.85) | 0.76 | 0.63 | 0.75 |
|  | Severe Sepsis | 0.87<br>(0.86, 0.88) | 0.80<br>(0.79, 0.81) | 0.77 | 0.65 | 0.72 |
|  | Septic Shock | 0.9992<br>(0.9991, 0.9994) | 0.963<br>(0.959, 0.968) | 0.94 | 0.86 | 0.82 |
| Sensitivity<br>(*Specificity fixed*<br>*near 0.80*) | Sepsis | 0.98<br>(0.96, 1.00) | 0.99<br>(0.97, 1.00) | 0.98 | 0.82 | 0.82 |
|  | Severe Sepsis | 0.996<br>(0.989, 1.000) | 1.00<br>(1.00, 1.00) | 0.98 | 0.90 | 0.81 |
|  | Septic Shock | 1.00<br>(1.00, 1.00) | 0.994<br>(0.992, 0.997) | 1.00 | 0.99 | 0.91 |
| Specificity<br>(*Sensitivity fixed*<br>*near 0.80*) | Sepsis | 0.95 (0.93, 0.97) | 0.75 (0.73, 0.77) | 0.72 | 0.32 | 0.51 |
|  | Severe Sepsis | 0.85 (0.84, 0.86) | 0.68 (0.62, 0.75) | 0.72 | 0.37 | 0.50 |
|  | Septic Shock | 0.9990<br>(0.9987, 0.9993) | 0.95<br>(0.94, 0.96) | 0.91 | 0.58 | 0.49 |

In addition to *InSight’s* ability to detect sepsis, severe sepsis, and septic shock, Figure 3A illustrates the ROC of severe sepsis detection and prediction four hours prior to severe sepsis onset. Even four hours in advance, the *InSight* severe sepsis AUROC is 0.85 (95% CI 0.79 - 0.91), which is significantly higher than the onset time SIRS result of 0.75 AUROC. Figure 3B summarizes *InSight’s* predictive advantage, using the severe sepsis gold standard, over MEWS, SOFA, and SIRS at the same time points in the hours leading up to onset. *InSight* maintains a high AUROC in the continuum up to four hours preceding severe sepsis onset. *InSight’s* predictions four hours in advance produce a sensitivity and specificity that are greater than the at-onset time sensitivity and specificity of each MEWS, SOFA, and SIRS (Table 2, Fig. 3B). In order to determine the generalizability of the algorithm to different settings, we tested InSight on additional patient data sets from four distinct hospitals. For severe sepsis detection at time of onset, *InSight* achieved AUROC over 0.92 on patients from Stanford, Oroville Hospital, Bakersfield Heart Health, and Cape Regional Medical Center (Table 3). ROC curves and comparisons to alternate sepsis classification systems on these datasets are presented in the data supplement (Supplementary Tables 5-8, Supplementary Figures 1 and 2). *InSight* AUROC values exceed those of the MEWS, SIRS, qSOFA and SOFA scores on the same datasets for severe sepsis detection at time of onset.

**Figure 3:**
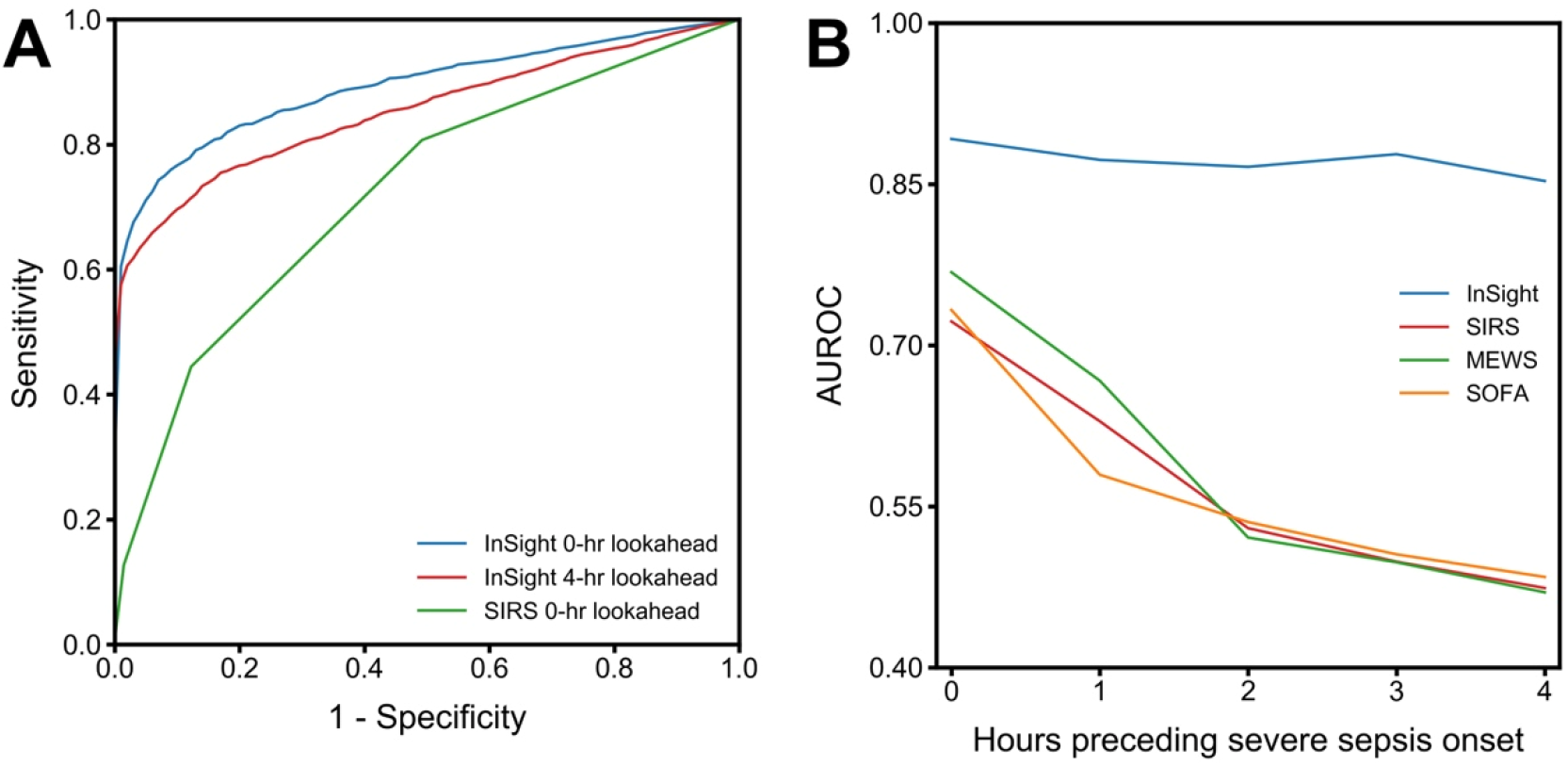
A) ROC detection (zero hour, blue) and prediction (four hour prior to onset, red) curves using *InSight* and ROC detection (zero hour, green) curve for SIRS, with the severe sepsis gold standard. B) Predictive performance of *InSight* and comparators, using the severe sepsis gold standard, as a function of time prior to onset.

**Table 3.** Algorithm performance for severe sepsis detection at time of onset. LR= Likelihood ratio.

|  | Stanford | Oroville | BHH | CRMC |
| --- | --- | --- | --- | --- |
| AUROC<br>(95% CI) | 0.924<br>(0.9202,0.9278) | 0.983<br>(0.9804, 0.9856) | 0.945<br>(0.921,0.969) | 0.960<br>(0.954, 0.966) |
| Sensitivity | 0.798 | 0.806 | 0.875 | 0.802 |
| Specificity | 0.901 | 0.989 | 0.940 | 0.946 |
| Accuracy | 0.900 | 0.971 | 0.963 | 0.931 |
| LR+ | 8.253 | 77.92 | 58.94 | 16.85 |
| LR- | 0.224 | 0.197 | 0.129 | 0.210 |

We ranked feature importance for the classifiers developed in this experiment, and determined that systolic blood pressure at the time of prediction was consistently the most important feature in making accurate model predictions. The relative importance of other features varied significantly based on the specific prediction task.

In our second set of experiments, we validated *InSight’s* performance in the presence of missing data. We tested *InSight’s* ability to detect severe sepsis at time of onset with various rates of data dropout. Table 4 presents the results of these experiments. After randomly deleting data from the test set with a probability of 0.10, *InSight’s* AUROC for severe sepsis detection is 0.82. Dropping approximately 60% of the test set measurements results in an AUROC of 0.75, demonstrating *InSight’s* robustness to missing data. Of note, the AUROC of *InSight* at 60% data dropout achieves slightly better performance than SIRS with no missing data. Further, our experiments on applying *InSight* to up- and down-sampled sets showed that AUROC was largest when the set was chosen such that around half the patients met the gold standard. Moving lower on prevalence from 50% down to 0%, the AUROC values were only slightly lower while they dropped steeply when moving higher on prevalence from 50% up to 100% (a clinically unrealistic range).

**Table 4:** *InSight’*s severe sepsis screening performance at time of onset in the presence of data sparsity, compared to SIRS with a full data complement.

|  | *InSight* |  |  |  |  | SIRS |
| --- | --- | --- | --- | --- | --- | --- |
| % Data Missing | 0% | 10% | 20% | 40% | 60% | 0% |
| AUROC | 0.90 | 0.82 | 0.79 | 0.76 | 0.75 | 0.72 |
| Sensitivity | 0.80 | 0.80 | 0.80 | 0.80 | 0.80 | 0.80 |
| Specificity | 0.84 | 0.66 | 0.57 | 0.50 | 0.49 | 0.51 |

## Transfer Learning

*InSight* is flexible by design, and can be easily trained on an appropriate retrospective data set before being applied to a new patient population. However, sufficient historical patient data is not always available for training on the target population. We evaluated *InSight*’s performance when trained on a mixture of the MIMIC-III data together with increasing amounts of UCSF training data, and then tested on a separate hold-out UCSF patient population using transfer learning. In Figure 4, we show that the performance of the algorithm improves as the fraction of UCSF target population data used in training increases.

**Figure 4.**
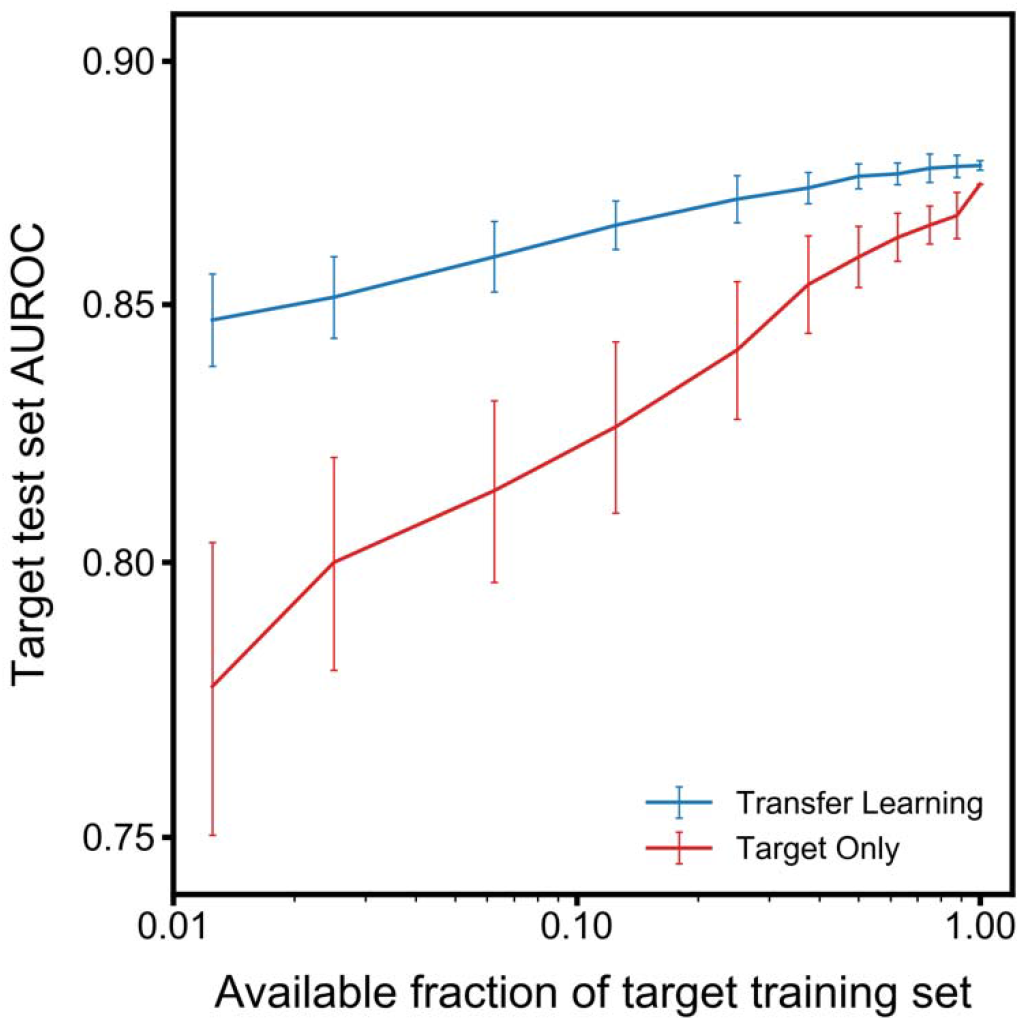
Learning curves (mean AUROC on the UCSF target data set) with increasing number of target training examples. Error bars represent the standard deviation. When data availability of the target set is low, target-only training exhibits lower AUROC values and high variability.

Feature importance was quite stable across transfer learning experiments, with systolic blood pressure measurements consistently playing an important role. Systolic blood pressure at two hours before onset, at time of onset, and at one hour before onset, in that order, were the most important features for accurate prediction in all tasks. Heart rate and diastolic blood pressure at time of onset were consistently the fourth and fifth most important features, though order of importance of the two features varied between tasks.

## Discussion

We have validated the machine learning algorithm, *InSight*, on the mixed-ward data of UCSF, which includes patients from the ED and floor units as well as the ICU, with varying types and frequencies of patient measurements. *InSight* outperformed commonly-used disease severity scores such as SIRS, MEWS, and SOFA for the screening of sepsis, severe sepsis, and septic shock (Figure 2). These results, shown in Table 2, confirm *InSight*’s strength in predicting these sepsis-related gold standard outcomes. The algorithm’s strong performance across the academic and community hospital data used in this study suggests potential strong performance in a variety of future clinical settings.

To the authors’ knowledge, *InSight* is first sepsis screening system to meet or exceed an AUROC of 0.90 using only vital sign inputs, on each of the sepsis gold standards evaluated in this study. Additionally, *InSight* provides predictive capabilities in advance of sepsis onset, aided by the analysis of trends and correlations between vital sign measurements. This advantage is apparent in the comparison with SIRS made in Figure 3A. Up to four hours prior to severe sepsis onset, *InSigh*t maintains a high AUROC above 0.85 (Figure 3). This advance warning of patients trending toward severe sepsis could extend the window for meaningful clinical intervention.

*InSight* uses only six common vital signs derived from a patient’s EHR to detect sepsis onset, as well as to predict those patients most at risk for developing sepsis. The decreased performance of *InSight* for recognition of severe sepsis relative to sepsis onset may be in part because the organ failure characteristic of severe sepsis is more easily recognizable through laboratory tests for organ function. Because we have not incorporated metabolic function panels in this validation of *InSight*, the detection of organ failure using only six common vital signs may be more difficult. In practice, *InSight* is adaptable to different inputs and is able to incorporate laboratory results as they become available. Inclusion of these results may well increase the performance of *InSight* for the detection and prediction of severe sepsis. However, in this work we have chosen to benchmark the performance of *InSight* using only six commonly measured vital signs. The ordering of metabolic panel laboratory tests are often predicated on clinician suspicion of severe sepsis, and therefore, early or developing cases may be missed. Additionally, because these vital sign inputs do not require time-dependent laboratory results or additional manual data entry, surveillance by *InSight* is frequent, and as a result, sepsis conditions are detected in a more timely manner. Minimal data requirements also lighten the burden of implementation in a clinical setting and broaden the potential clinical applications of *InSight*.

Although *InSight* uses only a handful of clinical variables, it maintains a high level of performance in experiments with randomly missing data. We demonstrate in Table 4 that for the detection of severe sepsis, even with up to 60% of randomized test patient data missing, *InSight* still achieves slightly better performance to SIRS calculated with complete data availability.

Additionally, we have investigated the customizability of *InSight* to local hospital demographics and measurements. The incorporation of site-specific data into the training set using transfer learning improves performance on test sets, over that of a training set comprised entirely of an independent population. This indicates that it may be possible to adequately train *InSight* for use in a new clinical setting, while still predominantly using existing retrospective data from other institutions. Further, the results of our up- and down-sampling experiments indicate that *InSight* is likely to only be slightly less effective (in AUROC terms) in settings with lower prevalence of sepsis, severe sepsis or septic shock, than UCSF or slightly more effective if the prevalence is higher than UCSF.

Our previous studies, performed on earlier versions of the model, have investigated *InSight* applied to individual sepsis standards such as the SIRS standard for sepsis [21], severe sepsis [22], and septic shock [23], on the MIMIC retrospective datasets. We have also developed a related algorithm to detect patient stability [24] and predict mortality [25, 26]. However, this study, which evaluates a significantly improved algorithm, is the first to apply *InSight* to all three standard sepsis definitions simultaneously, and to validate the algorithm on a mixed ward population, including ED, ICU and floor wards from UCSF. This study is also the first to use only six minimal vital signs, without utilizing a mental status evaluation such as Glasgow Coma Score, or even age, in the detection and prediction of those sepsis standards.

The separate models trained for each gold standard and prediction window in this study further demonstrate the potential clinical utility of machine learning methods. In addition to training on a specific patient population, machine learning methods can allow for the development of prediction models which are tailored to a hospital’s unique needs, data availability, and existing workflow practices. Any one of the models developed in this study could be independently deployed in a clinical setting; choice of model deployment would be contingent upon the needs of a particular hospital, and the expected tradeoff in performance for different model choices. Additionally, this study demonstrates the adaptability of the machine learning algorithm to an entirely new patient data set with markedly different demographics and outcomes through both site-specific retraining and transfer learning techniques.

## Limitations

While we incorporated data from multiple institutions, we cannot claim generalizability of our results to other populations on the basis of this study alone. However, we are aided by the minimality of data used to make predictions; because *InSight* requires only six of the most basic and widely-available clinical measurements, it is likely that it will perform similarly in other settings if vital sign data is available. The gold standard references we use to determine sepsis, severe sepsis and septic shock rely on ICD-9 codes from the hospital database; this standard potentially limits our ability to capture all septic patients in the dataset, should any have been undiagnosed or improperly recorded. The administrative coding procedures may vary by hospital and do not always precisely reproduce results from manual chart review for sepsis diagnosis, although ICD-9 codes have been previously validated for accuracy in the detection of severe sepsis [27]. The vital sign measurements abstracted from the EHR are basic measurements routinely collected from all patients regardless of diagnosis and independent of physician judgement, and therefore this input to *InSight* is not dependent on the time of clinical diagnosis. However, the ordering of laboratory tests is contingent on physician suspicion, and the timing of these inputs may reflect clinician judgement rather than true onset time, potentially limiting the accuracy of our analysis.

While the imputation and averaging performed before feature construction eliminated some information about sampling frequency, these methods do not remove all non-physiological information inherent to our system. Further, imputation of the most recently available past measurement may artificially alter the rate of the temporal changes in patient vital signs that we incorporate into feature vectors, which may in turn affect risk predictions. Averaging multiple patient measurements may similarly remove informative variation in vital signs.

It is important to note that we designed the study as a classification task rather than a time-to-event modeling experiment, because the former is significantly more common in the literature [28-31]. The alternative would not allow for the use of an established, standard set of performance metrics such as AUROC and specificity without custom modification, and would make it more difficult to compare the present study to prior work in the field. This study was conducted retrospectively, and so we are unable to make claims regarding performance in a prospective setting, which involves the interpretation and use of *InSight*’s predictions by clinicians. Additionally, our inclusion criteria requiring at least seven hours of patient data preceding sepsis onset also limits generalizability to a clinical setting where the predictor would receive data in real time. Algorithm performance in a clinical setting may reasonably be expected to be lower than its retrospective performance in this study. Finally, our random deletion of data is not necessarily representative of data scarcity as it would occur in clinical settings where the rate of missing measurements would depend on the standard rate of data collection, which can vary widely, especially between the emergency department, general ward, and intensive care units. We intend to evaluate these algorithms in prospective clinical studies in future work.

## Conclusions

We have validated the machine learning algorithm, *InSight*, in a multicenter study in a mixed-ward population from UCSF and an ICU population from BIDMC. *InSight* provides high sensitivity and specificity for the detection and prediction of sepsis, severe sepsis, and septic shock using the analysis of only six common vital signs taken from the electronic health record. *InSight* outperforms scoring systems in current use for the detection of sepsis, is robust to a significant amount of missing patient data, and can be customized to novel sites using a limited amount of site-specific data. Our results indicate that *InSight* outperforms tools currently used for sepsis detection and prediction, which may lead to improvements in sepsis-related patient outcomes.

## Acknowledgments

We acknowledge the assistance of Siddharth Gampa, Anna Lynn-Palevsky and Emily Huynh for editing contributions. We thank Dr. Hamid Mohamadlou and Dr. Thomas Desautels for contributions to the development of the machine learning algorithm *InSight*. We acknowledge Zirui Jiang for valuable computational assistance. We gratefully thank Matthew N. Fine, MD, Dr. Andrea McCoy, and Chris Maupin, RN for access to patient data sets.

## Author Statement

QM, JC, and RD conceived the described experiments. DS acquired the UCSF data. QM and YZ executed the experiments. QM, RD, JC, and MJ interpreted the results. QM, MJ, and JH wrote the manuscript. QM, RD, MJ, JH, JC, CB, DS, LS, UC, GF, and YK revised the manuscript, with assistance from Emily Huynh and Siddharth Gampa. All authors approved the version to be published and agree to be accountable for all aspects of the work in ensuring that questions related to the accuracy or integrity of any part of the work are appropriately investigated and resolved.

## Data Sharing

No data obtained from UCSF, Stanford, Oroville Hospital, Cape Regional Medical Center or Bakersfield Heart Hospital in this study can be shared or made available for open access. MIMIC-III is a publicly available database. Please visit https://mimic.physionet.org/ for information on using the MIMIC-III database.

